# How do dairy farmers wish their future farm?

**DOI:** 10.1101/777201

**Authors:** Anne-Catherine Dalcq, Thomas Dogot, Yves Beckers, Yves Brostaux, Eric Froidmont, Frédéric Vanwindekens, Hélène Soyeurt

**Author notes:** Corresponding author (ACD). Ideal future dairy farms to ensure revenue.

## Abstract

Dairy farming systems are evolving. This study presents dairy producers’ perceptions of their ideal future farm (**IFF**) to ensure revenue and attempts to determine the reasons for this choice, the environmental aspects related to this choice, the proximity between the current farm and the IFF and the requirements for reaching this IFF. Just before the end of the European milk quota, a total of 245 dairy producers answered a survey about the characteristics of their IFF and other socio-environmental-economic information. A multiple correspondence analysis (**MCA**) was carried out using seven characteristics of IFF (intensive *vs.* extensive, specialised vs. diversified, strongly *vs.* weakly based on new technologies, managed by a group of managers *vs.* an independent farmer, employed *vs.* familial workforce, local *vs.* global market, standard *vs.* quality-differentiated production). Based on the main contributors to the second dimension of MCA, this axis was defined as an IFF gradient between the local-based extensive (**LBE**) producers (26%) and the global-based intensive (**GBI**) producers (46%). The differences of IFF gradient between modalities of qualitative variables were estimated using generalised linear models. Pearson correlations were calculated between the scores on the IFF gradient and quantitative variables. Finally, frequencies for IFF characteristic and the corresponding characteristic for the current situation were calculated to determine the percentages of “unhappy” producers. Some reasons for the choice of IFF by the producers have been highlighted in this study. Environmental initiatives were more valued by LBE than GBI producers. Low similarity was observed between the current farm situation of the respondents and their IFF choice. LBE and GBI producers differed significantly regarding domains of formation (technical and bureaucratic vs transformation and diversification respectively) and paths of formation (non-market vs. market respectively). Two kinds of farming systems were considered by dairy producers and some socioeconomic and environmental components differed between them.

## Introduction

Food is a basic need. Working to provide food for themselves and their family was the task of everyone at the dawn of humanity. The progressive organisation of society has led to the appearance of “producers” who are responsible for producing food for more than just themselves and their family. Since World War II, public policies have been set up to increase food production [1]. These policies impacted the development of producers and their farms in the European Union. In the southern part of Belgium, the mean number of cows and the mean agricultural area per producer increased between 1980 and 2017 from 20 to 66 heads and from 25 to 71 hectares, respectively [2].

Producers are now facing great challenges to stay profitable. The price of the inputs (e.g., buildings, agricultural machinery, installations, feeding, veterinary care) of dairy production (**DP**) are increasing while the milk price shows great variability and its inflation is not similar to that observed for the inputs [3, 4]. Moreover, the European Union has decreased financial support to farmers [5]. On 1^st^ April 2015, the European Union removed the quota system which had managed the supply of DP [6]. This led to greater milk price volatility. Additionally, sanitary crises such as mad cow disease (bovine spongiform encephalopathy (BSE)) and the dioxine crisis, among others, have shocked consumers and led to new rules and regulations at European level and led to the creation of food security agencies in its countries. Moreover, these episodes modified consumers’ behaviours regarding their food purchases, they asked for more transparency and directed themselves towards organic food or local chains [7]. Besides the economic view, the impacts of farming on the environment have been noted and policies have been set up in the Common Agricultural Policy to solve these problems [4, 8].

In this context, the question often asked is what the future of dairy farming entails, how to remain profitable and more generally sustainable. Several authors have studied the evolution of dairy farming and the present dairy systems, finding trends that exist in the sector [3, 4, 9]. For instance, the project Mouve, funded by the French National Research Agency, studied the evolution of dairy farming systems in 6 dairy basins around the world. Their results gathered the publications of Napoleone *et al.* (2014) [9] and Havet *et al.* (2015) [3]. Moreover, authors have studied the future paths of development considered by dairy producers [10-13]. These studies were realised on the basis of data from 2001 to the beginning of 2013. These studies explored some reasons for these choices [10-14].

This study is innovative as it asks a different question to the other studies: what is the ideal future farm (**IFF**) to ensure revenue. Moreover, respondent producers were asked not to take into account their current farm. The data collection was conducted at the end of 2014 and the beginning of 2015. This was a particular context, just before the quota removal, when producers had this new perspective in mind and following two important milk crises associating low milk price due to a deregulation of EU milk production in 2009 and an increase of the cost of inputs in 2012. This research studied unprecedented reasons for this choice compared to what is present in the literature, to our knowledge, such as past events of the farms. Moreover, the present study explored the environmental and formational aspects linked to this IFF vision. The environmental aspect is of high importance at a time of increasing awareness of the impacts of agriculture and breeding on the environment. The formation aspect was studied to orientate university and other stakeholders of breeding improvement towards the domains needed by dairy producers. A comparison between the current farm and the IFF of the respondent was realised, and permitted the difference between the reality and the aspiration of the producers to be studied. More specifically, the goals of this study are to describe the perception of dairy producers about their ideal future dairy farming systems, and to quantify the proportion of producers desiring different dairy systems. By gathering different kinds of information, of which some are novel or a rarely present in the literature, this study also analyses the relationships between the dairy farming system desired by the producers and the reason(s) for this choice, their considerations about the environment and their needs to reach their goal of formation. Finally, we have ended this study by measuring the proximity between their current farming system and their IFF.

## Materials and methods

All editing and statistical analyses were carried out using SAS software (version 9.4., SAS Inst. Inc., Cary, NC, USA).

### Survey and IFF typology

A total of 245 Walloon dairy producers answered a survey between November 2014 and January 2015, the period just before the quota removal (1^st^ April 2015). Dairy producers answered questions at a time when they had this new perspective in mind. The sample set represented 6.1% of the dairy producers in Wallonia (about 4000 dairy producers in 2015 and 3500 in 2017 [2]). The density of dairy farms throughout Wallonia was well represented in the sample, with a higher answer rate in the provinces more populated with dairy farms. More answers were obtained in the east part of Belgium, where a higher density of dairy farms exists due to the grazing landscape that is particularly suitable for dairy production. Wallonia is a highly heterogeneous region with regard to soil and geological characteristics [15].

Dairy producers of the survey declared a mean of 79 cows and 86 hectares. Dairy production was the unique activity for 33% of them.

The entire survey was composed of 127 questions where the answers were decomposed into 498 qualitative and 44 quantitative variables. Concerning the perception of the IFF desired by the dairy producers to ensure revenue, 7 questions dealt with the following aspects: intensive *vs.* extensive production; specialised *vs.* diversified activity (or activities); farming strongly *vs.* weakly based on new technologies; farm managed by an independent farmer *vs*. a group of managers; family *vs*. employed workforce; providing production for local *vs*. global markets; and providing standard *vs*. differentiated quality production. The modality “No opinion” was available for each IFF question. Counts were calculated for all modalities of these seven questions.

To study the relationships between all modalities derived from the seven questions asked, a multiple correspondence analysis (**MCA**) was carried out as the variables were qualitative. For MCA, the eigenvalue of the dimensions generated, named principal inertia, is a biased measure of the part of information presented by a dimension [16]. Corrected inertia rates were calculated, as described by Benzécri (1979) [17], to quantify the correct part of information of a dimension.

Classes were established to study the distribution of producers along the dimensions of MCA. The interval between the 1% percentile and the 99% percentile of each dimension was divided equally into five classes. Then, the individuals per class were counted.

To exclude a group of producers with particular characteristics if necessary, cluster analysis was used on the scores of the individuals on the dimension of MCA, using the PROC CLUSTER procedure with the WARD method option.

### Characterisation of IFF choice

To describe the dairy producers in terms of their IFF, the scores on MCA dimensions were studied as a function of other variables extracted from the survey and distributed within several themes. These were the effect of past crises, problems encountered by the farmer, production factors, age of the farmer, breed of the cow, diversification of activities and alternative valorisation, regrouping between producers, consideration of mechanisation and robotisation on the farm, the reaction of the farmer to external factors, the considerations of farmers about environmental aspects, climatic hazard, ways to reach the ideal formation and field of formation. For qualitative variables, the scores of MCA dimensions were modelled using these variables as a fixed effect in a generalised linear model. Least squares means were estimated for the two-by-two comparisons using the Tukey test. The level of significance of those differences was assessed based on the *P*-value of the test. For quantitative variables, Pearson correlation coefficients were calculated between the scores of MCA dimensions and these variables. Their corresponding *P*-values were estimated to observe if the correlation values were significantly different from 0.

To observe if dairy producers presented the farming characteristics considered as ideal at the moment of survey, absolute frequencies were calculated as a function of each ideal future farm characteristic and the corresponding characteristic for the current situation (Table 1). Moreover, the percentage of “unhappy” producers was calculated as the ratio between the producers not currently in the situation that they consider as ideal and the total number of producers.

**Table 1.**
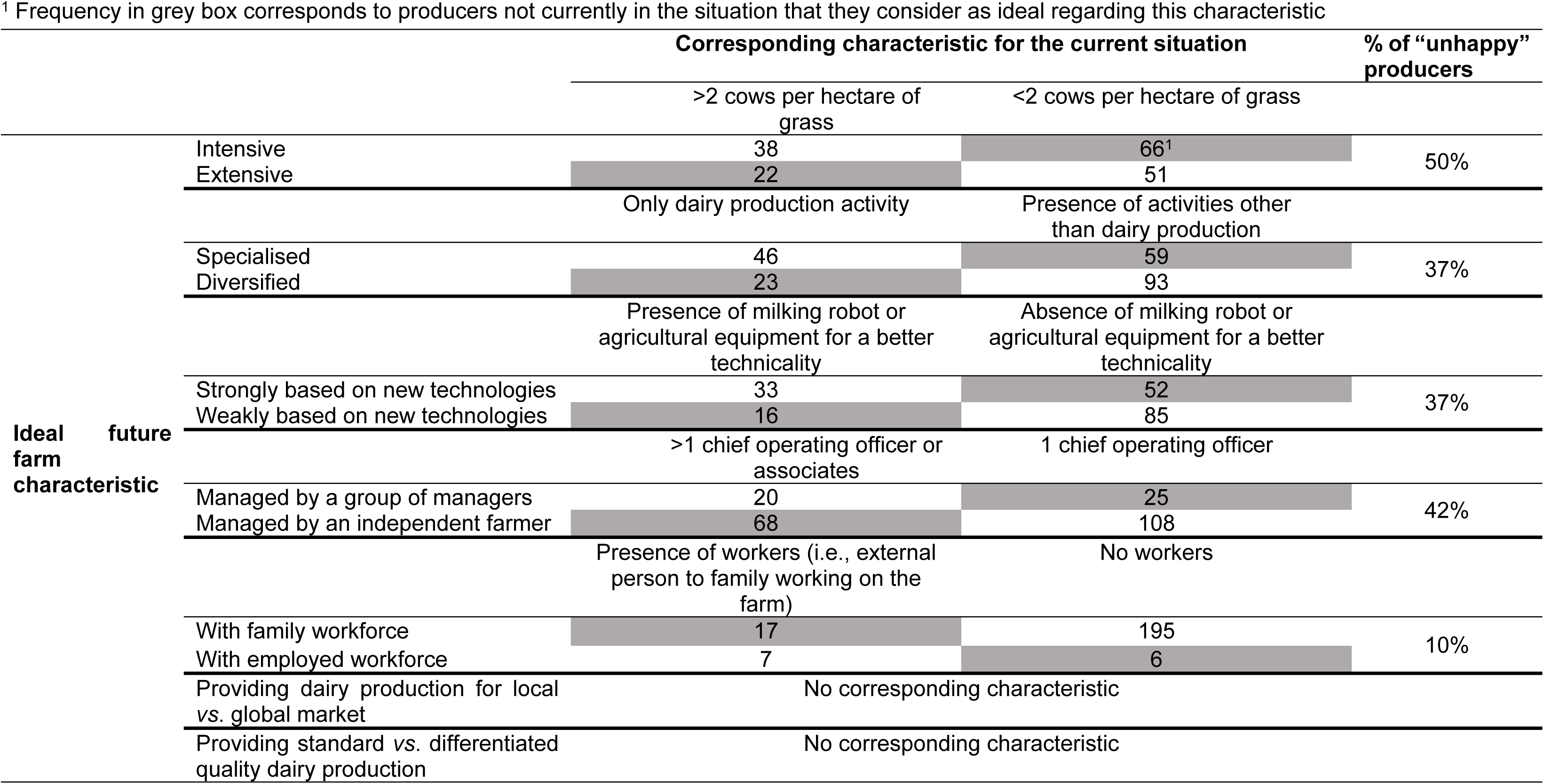
Absolute frequency of producers as a function of their answer to the ideal future farm characteristic and the corresponding characteristic for the current situation and percentage of “unhappy” producers (i.e., percentage of producers not currently in the situation that they consider as ideal)

## Results and discussion

### Contrasted opinions of Walloon dairy farmers about the ideal future farm

As mentioned previously, the first aim of this study was to highlight the perceptions of Walloon dairy producers about their ideal farm, just before the end of the milk quota. This was done through the answers to 7 questions. Table 2 shows the frequency for each modality of those questions.

**Table 2.** Percentages of the seven questions about the ideal future farm.

| Question | Proposition | Percentage (%) |
| --- | --- | --- |
| Intensive vs. extensive | Intensive | 43 |
|  | Extensive | 30 |
|  | No opinion | 27 |
| Specialised vs. diversified | Specialised | 43 |
|  | Diversified | 47 |
|  | No opinion | 10 |
| Strongly vs. weakly based on new technologies | Strongly | 35 |
|  | Weakly | 41 |
|  | No opinion | 24 |
| Managed by an independent farmer vs. a group of managers | Independent farmer | 72 |
|  | Group of managers | 18 |
|  | No opinion | 10 |
| Family vs. employed workforce | Family | 87 |
|  | Employed | 5 |
|  | No opinion | 8 |
| Providing dairy production for local vs. global market | Global | 43 |
|  | Local | 32 |
|  | No opinion | 25 |
| Providing standard vs. differentiated quality dairy production | Standard | 38 |
|  | Differentiated quality | 45 |
|  | No opinion | 17 |

Contrasted opinions of dairy farmers were observed for almost all questions except for the type of management and the kind of workforce: 71.84% of the respondents wanted an independent farmer management, and 86.53% focused on a family workforce (Table 2). These results highlight a will in the southern part of Belgium to maintain the traditional form of work organisation in the future, with family workforce and one director of operations. More globally in the world, dairy farms are still mostly owned and managed by a family structure, whatever the degree of development of the country [18, 19]. The choice of producers to work by themselves and not to deal with workers (i.e., an external person to the family employed on the farm) was noted in other studies. For example, in Spain Gonzalez and Gomez (2001) [20] observed, when asking 3,370 farmers for their definition of a farmer, that more than half of them chose labourer and 12% chose businessman.

From Table 2, it is remarkable to note that the highest percentages of abstention were observed for the questions about intensive *vs*. extensive, strongly *vs*. weakly based on new technologies, and providing DP for local *vs*. global markets. These results showed that a quite significant proportion of the respondents did not take a position on these directions for the evolution of dairy farms.

### Dualisation of ideal future farm aspirations

To study the relationships between the answers given by the respondents to all questions about IFF, a MCA was performed as the related variables were qualitative (Table 2). The percentage of principal inertia of the dimensions 1 and 2 of MCA were 16.75% and 12.38%, respectively (Fig 1). The value of corrected inertia for the two first dimensions reached 72.7% and 21.5% respectively, gathering almost 95% of the information.

**Fig 1.**
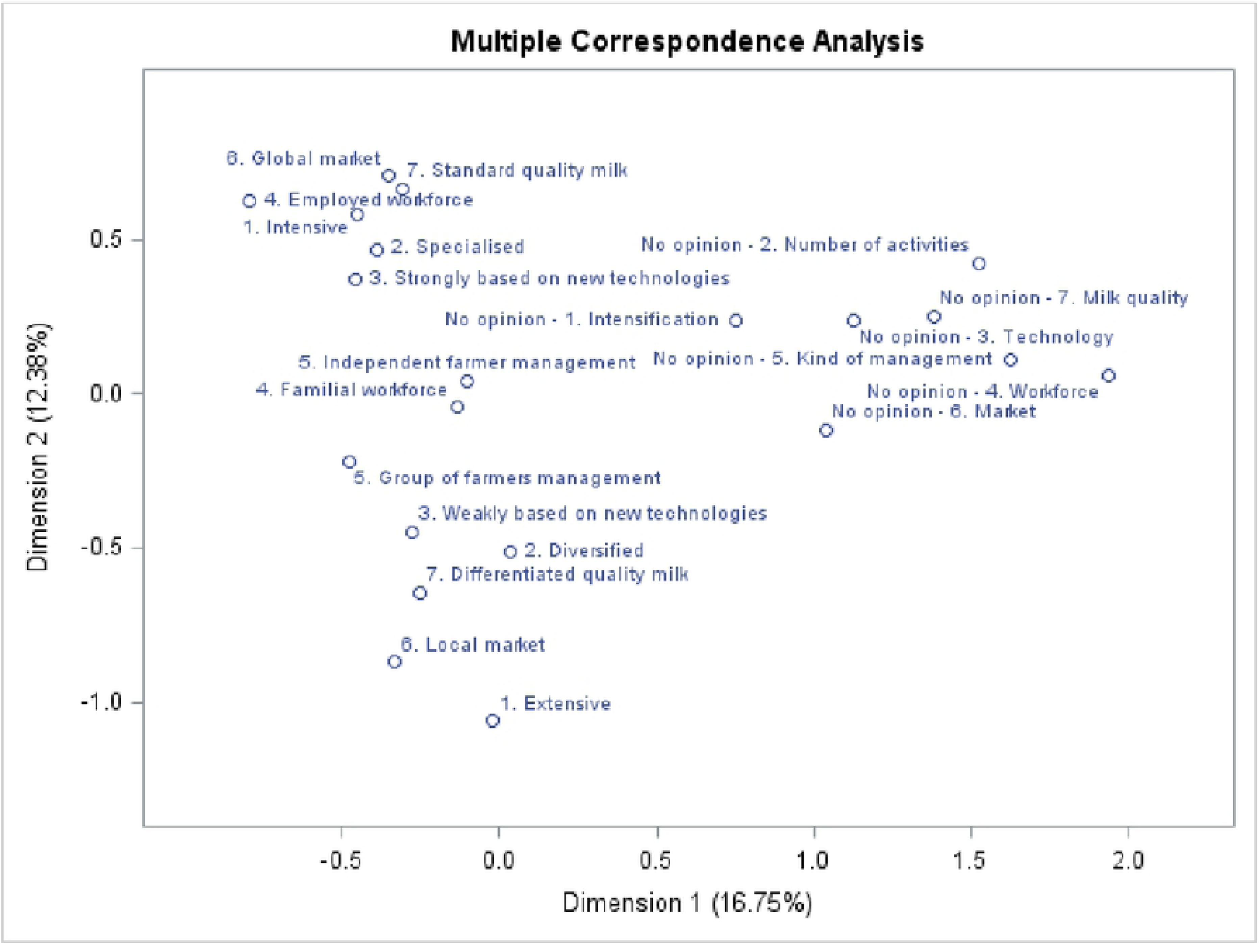
Representation of the modalities in the Multiple Correspondence Analysis first factorial plan. Values of principal inertia reached 16.75% and 12.38%. Values of corrected inertia reached 72.7% and 21.5%.

The first dimension permitted differentiation between the producers who did not give their opinion concerning characteristics of IFF and the producers who did (Fig 1). This represented 15% and 85% of the dataset respectively. The “no opinion” producers group (N = 38) were removed from the analysis to avoid potential bias coming from farmers who did not have a clear vision of their IFF.

The second dimension of the MCA was the most interesting for highlighting the wishes of dairy farmers about their IFF, for those who took position on this question. This axis showed a gradation of question modalities and proximity between several characteristics. This dimension led to the identification of two extreme tendencies (Fig 1); the modalities of familial workforce, independent farmer management and management by a group of farmers were near to zero on this axis (Fig 1). This means that the small proportion of producers supporting group management was distributed between the two extreme tendencies observed. The position of the modalities of familial workforce and independent farmer at the middle of the second dimension illustrated the fact that these modalities were chosen by producers from the two tendencies identified. The small proportion of producers choosing an employed workforce was positioned on the top of the second dimension (Fig 1).

The first tendency, related to high scores on the second MCA dimension, corresponds to IFF with the following characteristics: global market, standard milk, intensive system, employed workforce, specialised and strongly based on new technologies. Other authors have observed the same relations. From a trial of 458 French dairy farms, Hostiou *et al.* (2015) [21] highlighted a profile of farmers which simultaneously gathered high equipment, intensification and workers. From a trial of 3,370 producers of all sectors in Spain, Gonzalez and Gomez Benito (2001) [20] collated the characteristics of large holdings, market-orientated farming and management of workers. Cournut *et al.* (2010) [22] highlighted different ways of evolving dairy farming in France, characterised by workers, mechanisation and high equipment. This tendency in dairy farming systems is explained by the evolution of the dairy system [4]. The increased competition in the dairy market caused by the creation of the open European market as well as the wish of consumers to have structures that gather all the food supplies in one place (i.e., a supermarket) led to the concentration of dairy processing in some big firms [9]. These firms were better placed to develop because they could control their collection costs, benefit from scale economies and were able to deliver to supermarkets with regularity in quantity and with a standard quality [7]. This state and the world market have conditioned a milk price for the producers. Increasing the production, thanks to more cows or higher productivity, is a possible way to stay profitable, considering the undergone milk price [3, 9]. To achieve profitability, an elevated production of milk per cow and an increase of cows on the farm are reached [9]. Moreover, this increase in milk production at farm level was also forced by the orientated production Common Agricultural Policy (**CAP**) primes, although CAP has limited help to the dairy sector. Therefore, all of these characteristics intensify the dairy farming system. Intensification was defined by Garcia-Martinez *et al.* (2009) [23] as the maximisation of the rarest factor, traditionally the agricultural area. The increase in DP per unit of agricultural area was possible thanks to intensive production of forage and purchase of inputs, produced where production costs were the lowest, to balance the ration, to increase the production per cow, or the number of cows reared on a hectare of agricultural area and therefore DP per unit of agricultural area at the level of the farm [7, 9]. This intensification led to more specialised farms with more dairy cows and their entire workforce directed to this specialisation [7]. The enlargement of farms required a higher work rate; this was surmounted thanks to equipment and new technologies and thanks to more human workforce: collective organization, subcontracting to private firms and also employment of workers [7].

The second tendency, contradictory to the first tendency, was characterised by high negative scores on the second MCA dimension. This axis was represented by the following modalities: weakly based on new technologies, diversified, differentiated quality milk, local market and extensive system (Fig 1). This reflects another form of dairy farming. This form is favoured by a constant increase in input prices, combined with a growing demand of consumers to have high quality and local based products [7]. Dairy producers choose to work with greater self-sufficiency to be less dependent on the undergone input prices [7]. The “localisation” of the production demanded by consumers was executed thanks to this more local-produced forage and fewer inputs from outside [3]. This return to self-sufficiency led to more extensive farming [3]. The production induced was also often quality-differentiated and dedicated to local markets [7]. Cournut *et al.* (2012) [7] showed in their study that this kind of dairy farming is chosen by a minority of farms, which are still diversified.

This gradation with two kinds of models at the extremities of the second MCA dimension was also described in other studies [3, 4, 7, 9, 24-26]. They were named globalisation *vs*. territorialisation by Cournut *et al.* (2012) [7], or globalisation *vs*. localisation by Napoleone *et al.* (2014) [9]. Lebacq (2015) [4] identified a “dualisation of dairy farming systems between ‘a mainstream model’ focusing on an increasing farm size, production intensity and specialisation and alternative models involving initiatives deviating from this trend and constituting niche developments (niches = minor elements, hardly sustainable against the mainstream model)”.

Thanks to a survey answered by 180 producers of all sectors in 2007 in France, concerning the evolution of their farms and their aspirations, Dockes *et al.* (2007) [25] also highlighted a major tendency towards the enlargement, professionalisation and specialisation of farms, but those authors also mentioned that other farms wanted to develop diversified structures, orientated towards the requests of society, processing and farm accommodation. The present study showed this dualisation but also quantified these two tendencies: 46% *vs*. 26% of producers having high positive and high negative scores respectively on the second dimension. Verhees *et al.* (2018) [13] quantified producers as a function of their strategies of development, but solely regarding specialisation *vs*. diversification of their activity, 54.3% *vs* 15.1% respectively.

### Relationships between ideal future farms and reasons, environmental considerations and formations

To study the relationships between the different IFF and other interesting technico-economic information, the second dimension was considered as a gradient (**IFFg**) interpreted at the extremities as global-based intensive producers (GBI: high positive scores) and local-based extensive producers (LBE: high negative scores). The choice to work with a gradient rather than a clear separation of the two tendencies was motivated by the will to represent all the intermediaries of the IFF of dairy producers. The mean of the scores of the second MCA dimension was –0.012 with a SD of 0.053. Minimal and maximal values were –1.09 and 0.92, respectively.

Based on the interpretation of IFFg, a significant negative correlation indicates a higher relationship with the dairy producers desiring a LBE model. By opposition, a significant positive correlation means a higher link with the dairy producers desiring a GBI model. Tables 3, 5 and 6 give the results of generalised linear models where the qualitative variables were introduced separately as a fixed effect in the model. Significantly lower estimates of IFFg for a specific modality of the considered qualitative variable depicts a tendency of producers desiring a LBE model to choose this modality, while significantly higher estimates of IFFg means a tendency of producers wanting a GBI model to choose this modality. The following paragraphs will summarise the potential reasons driving the choice of IFF made by the Walloon dairy farmers.

**Table 3.**
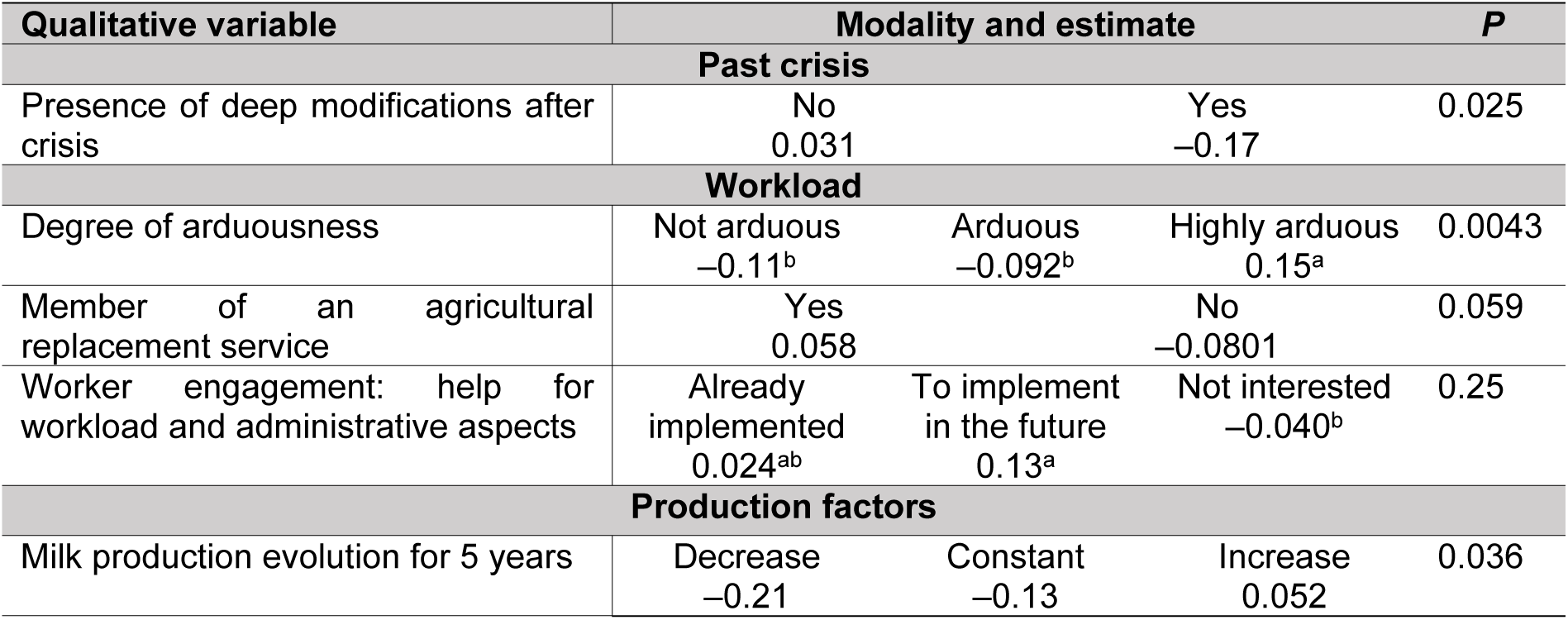
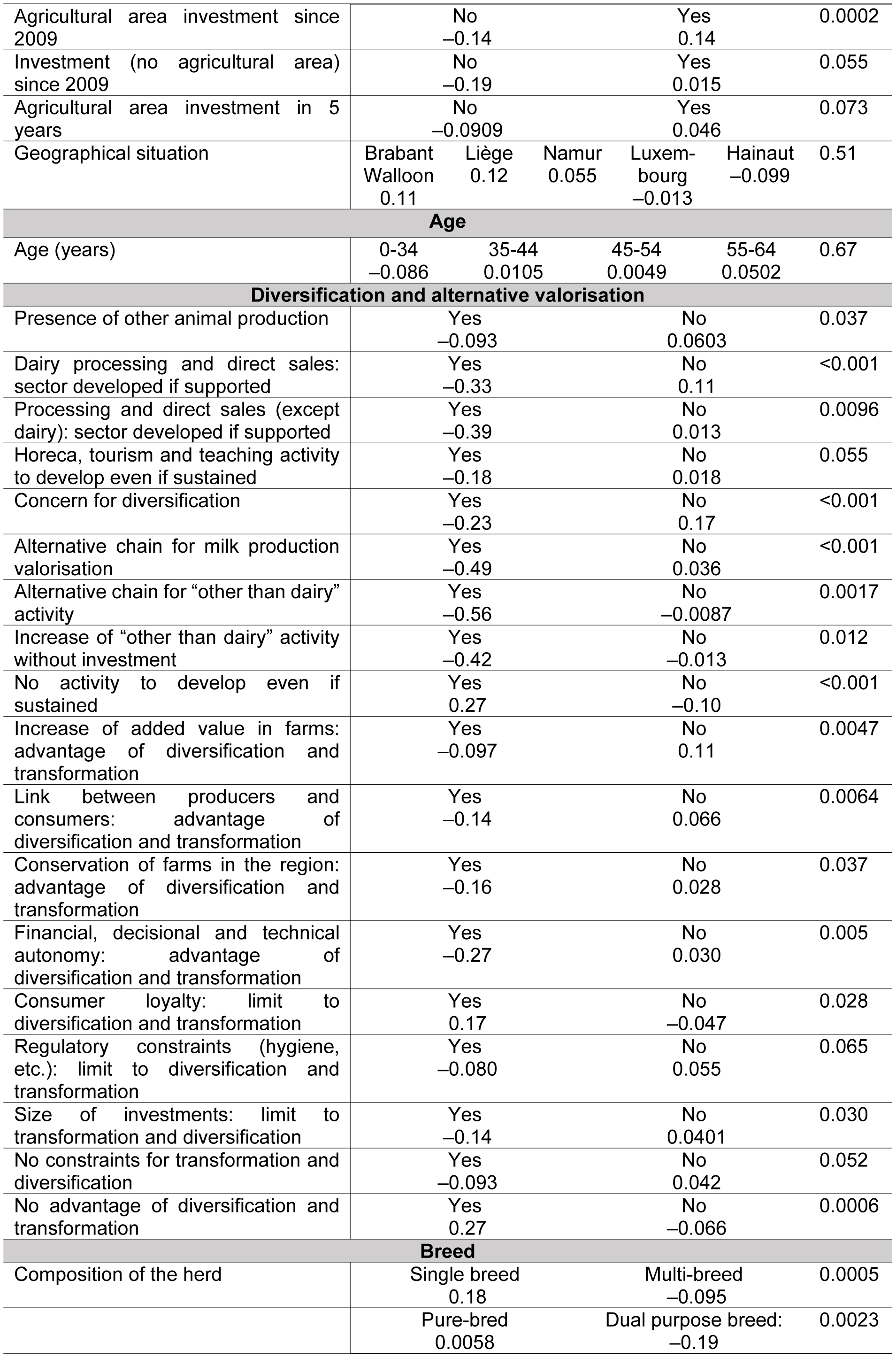
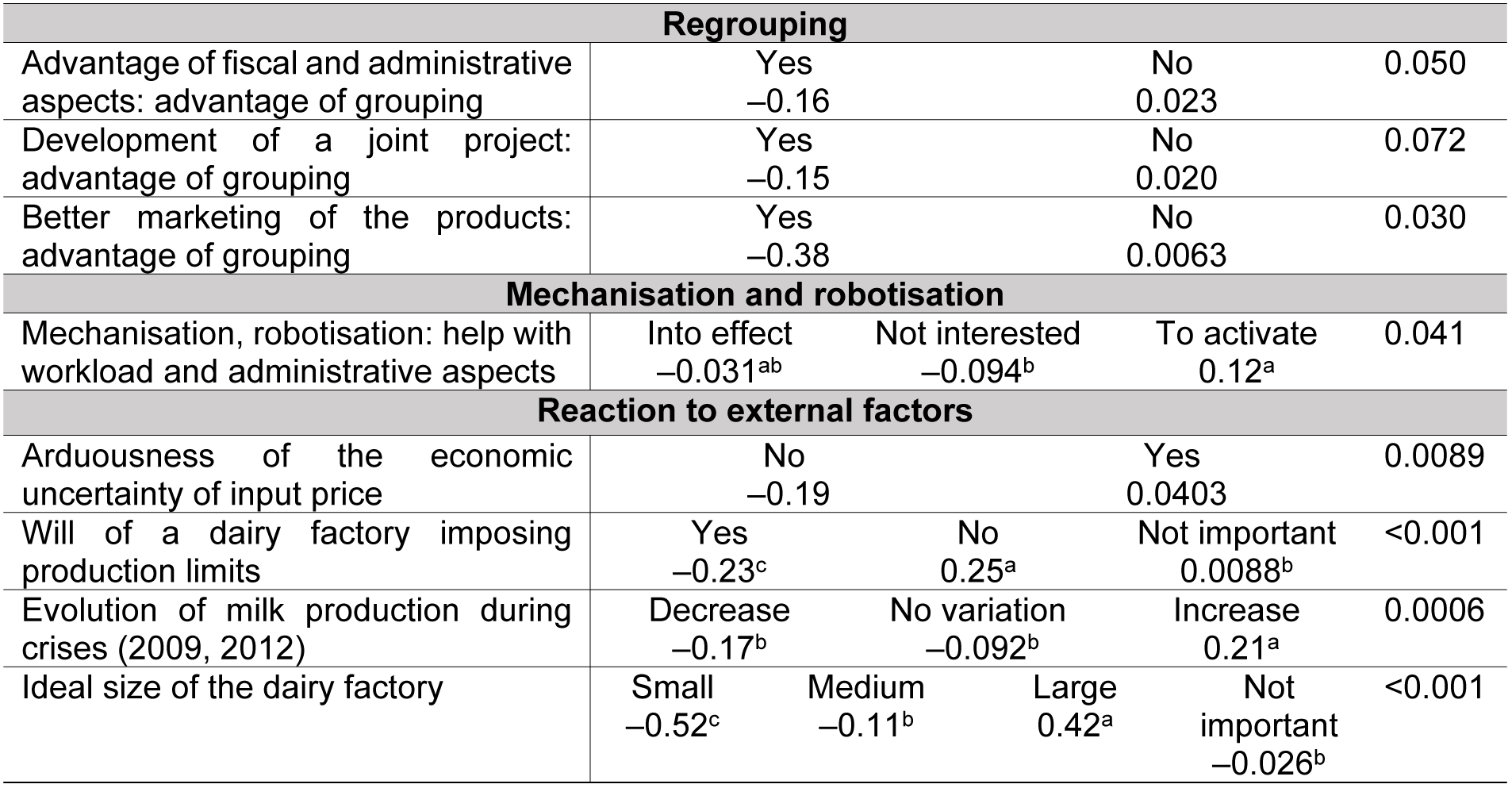
Value and level of significance of the difference in the ideal future farm gradient as a function of modalities of qualitative variables: reasons.

### Reasons

#### Effect of past crisis on perceptions of the ideal future farm

The producers that were impacted by past crises wished more for a LBE model (estimate = –0.17, Table 3). This could be related to the suffering involved in the crisis and the wish to apply solutions in order to not repeat this situation: revenue from diversified activities, other outlets for the milk production sold (i.e., local market characteristic) and/or self-sufficiency to be less dependent on purchased feed (i.e., extensive farm characteristic). This is in agreement with a past finding [27]. We observed a decrease in intensification in 2012 which was the year of a dairy economic crisis mainly related to an increase in the price of inputs.

#### Workload

Workload seems to be less bearable for producers desiring a GBI model (estimate = 0.15, Table 3; R _constraint workforce_ = 0.22, *P =* 0.002). Producers wishing for a GBI model were also nearly significantly more likely to be members of an agricultural replacement service (estimate = 0.058, Table 3) and showed a tendency to be more interested in employment of workers (estimate = 0.13, *P* _worker engagement to implement *vs.* not interested_= 0.11, Table 3). The choice of GBI model could be explained by this current workload, involving the need for an increase of revenue. So, the solution considered could be higher milk production and the breeding of more cows rather than diversification of activities and self-valorisation activity, the development of which requires a lot of time. Samson *et al.* (2016) [14] confirmed this in the Netherlands by highlighting a nearly significant effect of labour productivity on the DP increase strategy.

#### Production factors

The size of agricultural area, the milk delivery quota, the number of cows and the percentage of corn silage currently observed in the farming system showed significant and positive correlations with IFFg (R = 0.15, 0.36, 0.18 and 0.24; *P =* 0.033, <0.001, 0.0099, 0.0002 respectively). So, dairy producers choose their IFF partly as a function of their current production factors. This is expected as a higher number of hectares, cows and litres means a higher capacity of the dairy installation, of the material and so the possibility of a more preponderant dairy activity. The higher percentage of corn silage also reflected the possibility to seed corn silage, allowing the intensification of production as required within a GBI model. Similar relationships between characteristics of the farm and current or desired models of farming were observed by others. For Central and Eastern Europe, Verhees *et al.* (2018) [13] showed that land was the most important factor in developing a specific farming strategy. In France, Hostiou *et al.* (2015) [21] observed that intensified farms with higher technology equipment sometimes employed more workers, and were the farms with significantly higher agricultural area, percentage of corn silage, number of cows and milk quota. In the Netherlands, Samson *et al.* (2016) [14] showed that production intensity, number of cows, modernity of technology and availability of land were important factors in DP increase strategies.

In contrast, producers with lower production factors can consider more hardly enlargement and therefore think differently about the enhancement of their revenue: better valorisation of quality differentiated milk, other activities on the farm, self valorisation, the LBE model. Samson *et al.* (2016) [14] showed that lower stable capacity varies inversely to a DP increase strategy, which is rather a GBI tendency.

The findings of the current study, as confirmed by previous researchers, showed that producers work within a tightly constrained and regulated environment limiting their ability to determine the future of their farm according to their personal desires. This statement was also concluded by Mc Elwee *et al.* (2006) [28] and Methorst *et al.* (2017b) [29]. In the Netherlands, Keizer and Emvalomatis (2014) [30] and Groeneveld *et al.* (2016) [31] showed that bigger farms are more likely to increase than other farms.

However, based on the quite low values of the correlations obtained between the agricultural area and the number of cows, we can consider that this situation must be nuanced and that the IFF chosen also depends on the ways of thinking of the dairy producer, not taking into account the current situation of his farm. This statement is reinforced by the fact that the correlation of percentage of meadow with IFFg was not significantly different to 0 (R = –0.097, *P* > 0.1). Also, the impact of the provinces of the Walloon Region, which present different geographical and soil characteristics, on IFFg were not significantly different (*P* = 0.51, Table 3).

Moreover the significant relations between IFFg and milk production evolution for five years (Table 3; R _quantity of milk variation_= 0.30, *P* < 0.001), investment for and in five years (Table 3) support the assumption that the IFF chosen depends greatly on the mentality of the producers.

In their study, Methorst *et al.* (2017a) [11] proved the heterogeneity of farm developments of producers facing the same socio-material context, showing the importance of the mentality of the producers in their decisions. Authors speak about shared values, norms, ways they see themselves or would like to be seen by producers, views, capacities and their perceptions of opportunities and any room for manoeuvre, skills, motives, entrepreneurship, goals and strategies [10, 11, 14, 29, 32] as factors which influence the farm development. Samson *et al.* (2016) [14] discussed experimental economics, which are economics where psychology and biology, which explain human behaviours, are added to better explain the development of enterprises. The consideration of more than just economic aspects permits them to reduce the error of their model for predicting DP increase strategies [14].

#### Age

Age of the producer seems not to condition the desired IFF (Table 3). An IFF could be chosen because of either the new ideas of young producers or the experience of older producers. If mentality seems to influence IFF choice, it is not linked to age. The two kinds of IFF could be an answer to both innovation and problems encountered during a long career. Samson *et al.* (2016) [14] also studied age as a reflection of the farmers’ values, goals and strategies, and showed no relationship with DP increase, which is rather a GBI characteristic. On the contrary, on the basis of data from 11 countries of the European Union, Weltin *et al.* (2017) [12] observed an effect of age on the tendency towards diversification, which is rather a LBE tendency.

#### Diversification and alternative valorisation

The results obtained in this study showed a link between the diversification mentality and the choice of LBE model. Significant negative estimates or correlations were observed for the following variables related to diversification: the presence of other animal production (estimate = –0.093, Table 3); the direct selling milk quota (R = –0.17, *P =* 0.016); dairy or no dairy processing and direct sales (estimates = –0.33 and –0.39, Table 3); the development of HORECA activities, tourism and teaching (estimate = –0.18, Table 3); the concern for diversification (estimate = –0.23, Table 3); alternative chain for milk and other than milk production valorisation (estimates = –0.49 and –0.56, Table 3) and the increase of “other than dairy” activity without investment (estimate = –0.42, Table 3). Conversely, producers desiring a GBI model were more likely to choose the item “no activity to develop if supported”, suggesting the unique principal activity way of thinking of producers aiming for a GBI model (estimate = 0.27, Table 3). Samson *et al.* (2016) [14] confirmed this tendency and showed that the presence of diversified activities evolved inversely to the increase of milk production. In this study, we observed potential explanations to support to this fact. Producers wishing for a LBE model considered self-valorisation and diversification as solutions to the current situation to enhance revenue due to the creation of added value (estimate = –0.097, Table 3). They thought that diversification and transformation allowed financial, decisional and technical autonomy (estimate = –0.27, Table 3) and were confident in consumer loyalty (estimate = –0.047, Table 3). They considered relations with consumers as an opportunity and not a threat, unlike producers desiring a GBI model (estimate = 0.17, Table 3). One reason GBI model producers gave against self-valorisation and diversification seemed to be the lack of trust in consumers and therefore the outlets. They frequently saw no advantage to self-valorisation and diversification (estimate = 0.27, Table 3). The relation to the consumer was also studied by Verhees *et al.* (2018) [13]. They observed that consumer orientation was more often declared as an opportunity to the profiles of producers considering strategies similar to LBE. The positive impact of diversified activities on autonomy was also shown by Bergevoet *et al.* (2004) [10]. They mentioned that proponents of the “extra source of income” model (closest to the LBE model) were more able to declare that they can increase the sales-price of their milk. Producers wishing for a LBE model were also likely to find no constraints to transformation and diversification (estimate = –0.093, Table 3). The only limits to diversification and transformation highlighted by producers wanting a LBE model were regulatory constraints (estimate = –0.080, Table 3) and the size of investments (estimate = –0.14, Table 3). As a consequence of these considerations, producers wanting a LBE model felt that they were more able to meet society’s expectations regarding local and artisanal products (R = –0.22, *P =* 0.0016) and the desire for a familial structure (R = –0.12; *P =* 0.084).

#### Breed to produce milk

Producers wanting a LBE model are more open to breeding a dual-purpose herd (estimate = –0.19, Table 3), which permits them to diversify their production: milk and meat. Producers wishing for a GBI model target a single, more specialised breed (estimate = 0.18, Table 3) which could offer more homogeneous management of the herd. The link between mentality, observed through the choice of breed(s), and the choice of IFF is once more highlighted.

#### Regrouping

Producers tending towards the LBE model were more likely to promote regrouping for its advantages regarding fiscal and administrative aspects, the development of a joint project and the marketing of the products (estimates = –0.16; – 0.15; –0.38, Table 3). The importance of mentality for the choice of IFF has been shown. A mentality of cooperation, as a solution to enhance their quality of life and revenue, tends to be shared between producers desiring a LBE model.

#### Mechanisation and robotisation

A “pro-technology” mentality of the producers tending towards the GBI model was observed (estimate = 0.12, Table 3). It can be assumed that the solution considered by them is to keep the same activity or increase it with help from machines. In southern France, Dufour *et al.* (2007) [33] observed the propensity of farmers with workers, close to the GBI model, to prioritise investment in equipment. Verhees *et al.* (2018) [13] observed that better management, including new technologies, was more cited as an objective for strategy profiles of producers that were more similar to the GBI than LBE models.

#### Reaction to external factors

Reactions of dairy producers to factors external to their decision-making power tend to be different as a function of their choice of IFF, showing once more a different mentality of the producers. Producers wanting a LBE model tend to show themselves to be more independent from the external economic actors: from the input producing companies (estimate = –0.19, Table 3) and from the market and the factories, rejecting contracts which would link them to it (R = –0.13, Table 4). When their opinion about dairy factories was surveyed, producers desiring a LBE model preferred small or medium units with production limits (estimates = –0.52; –0.11; – 0.23, Table 3), as before, which means regulation of the dairy offerings on the market. Producers wishing for a GBI model direct themselves to big units of processing without production limits (estimates = 0.42; 0.25, Table 3) and so more turned towards world markets. They recognise the freedom in regarding DP as an asset of quota removal (R = 0.23, Table 4).

**Table 4.**
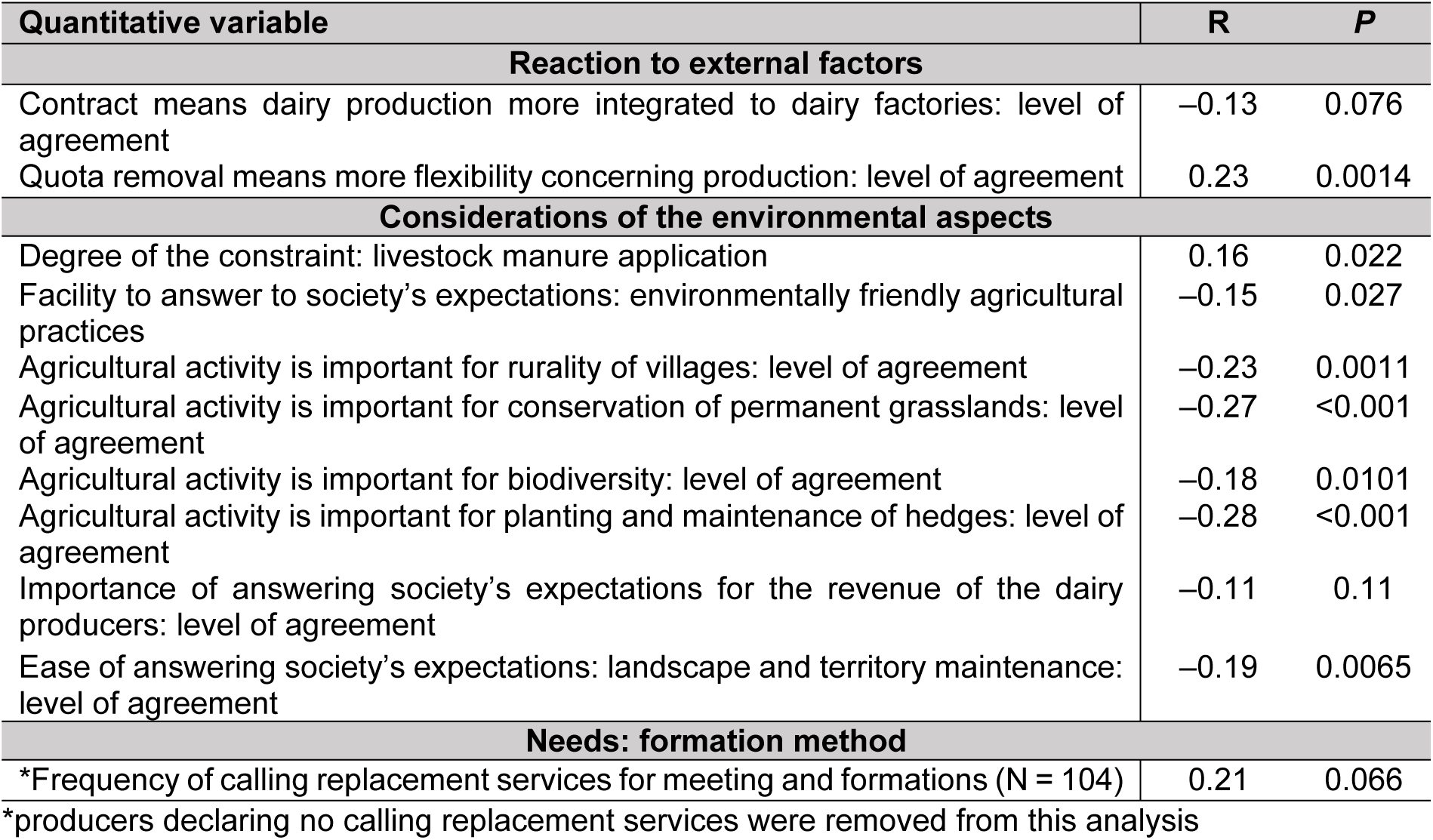
Correlations (R) between the ideal future farm gradient and quantitative variables.

The reaction regarding the quantity of production was not similar during a crisis, producers wanting a LBE model tended to maintain or decrease their production (estimates = –0.17; –0.092, Table 3), whereas producers desiring a GBI model tended to increase production (estimate = 0.21, Table 3). The latter wanted to keep revenues constant with more litres produced when the price decreased, while the others controlled or decreased production when the gross margin per litre decreased. It can be due to a deliberate choice to decrease the milk production or the decision to decrease the variables costs causing a decrease ok milk production. These results can express a fear of producers tending toward the LBE model in considering world markets, contrary to producers tending towards the GBI model who have decided to work with this kind of market. Verhees *et al.* (2018) [13] observed that producers projecting strategies similar to the LBE model consider the market more as a threat than producers projecting strategies similar to the GBI model. Hansson *et al.* (2010) [34] and Weltin *et al.* (2017) [12] explained that this uncertainty and risk perception can explain the choice of diversification, which is a part of the strategy of the LBE model.

Couzy and Dockès (2008) [5] demonstrated different profiles of farmers and observed the entrepreneurship mentality of each one, which highlights similar tendencies to those presented here. Several profiles showed strong entrepreneurship but which was expressed differently to here. A category of farmers showed entrepreneurship by their wish for autonomy of decision in their management; they will keep a working approach close to the conventional one but with a modernist vision, always adapting to the market. They want to keep freedom in the classical framework. Another category of farmers showed entrepreneurship by their wish to develop an original idea, away from preexisting systems, a project in line with their conviction to be freer from the existing system.

Samson *et al.* (2016) [14] and Methorst *et al.* (2017a) [11] reported that decisions of producers cannot be reduced to only economic aspects: this includes policies and market conditions but also their way of thinking about them.

### Consideration of environmental aspects

The environmental aspects related to the desired IFF is now studied as awareness of the environmental impact of breeding has become an important issue of our time.

Producers tending toward the GBI model seemed to work with a higher livestock manure application pressure (R = 0.16, Table 4) and therefore are already more to work in an intensified dairy system, which can impact the environment. Samson *et al.* (2016) [14] showed a tendency toward manure production surplus by producers with increasing DP, which is rather a GBI characteristic.

Results of practices that are in accordance with the environment: measurement of the grass height, forage mixture with leguminous plants, use of a field notebook (estimates = –0.27; –0.11; –0.074, Table 5) showed a stronger interest from producers wanting a LBE model.

**Table 5.** Value and level of significance for the difference in the ideal future farm gradient as a function of modalities of qualitative variables: environmental aspects.

| Qualitative variable | Modality and estimate |  | P |
| --- | --- | --- | --- |
| Considerations of the environmental aspects |  |  |  |
| Measurement of the grass height: optimisation practice | Yes<br>−0.27 | No<br>0.0083 | 0.059 |
| Forage mixture with leguminous plants: optimisation practice | Yes<br>−0.11 | No<br>0.083 | 0.0088 |
| Field notebook: optimisation practice | Yes<br>−0.074 | No<br>0.061 | 0.065 |
| Climatic hazard |  |  |  |
| Increase of concentrate distribution: strategy to confront climatic hazards | Yes<br>0.22 | No<br>−0.036 | 0.036 |
| Decrease of the herd: strategy to confront climatic hazards | Yes<br>−0.25 | No<br>0.014 | 0.037 |
| Food self-sufficiency: cause for maintaining constant or decreased milk production | Yes<br>−0.17 | No<br>0.0073 | 0.14 |

Besides these, all the significant negative correlations between IFFg and the levels of agreement with an agricultural area are important for the rurality of villages (R = –0.23, Table 4), for conservation of permanent grasslands (R = –0.27, Table 4), for biodiversity (R = –0.18, Table 4) and for hedges (R = –0.28, Table 4) showed the importance of the environment in the dairy activity of producers wanting a LBE model. It can be assumed that both LBE producers and GBI producers have concern for the environment but in a different way. These results showed that LBE producers are more willing to employ the benefits of ecosystem services, which is observable by these results. Moreover, they found it easy to realise environmentally friendly agricultural practices, as asked for by society (R = –0.15, Table 4) and which are important to answer to society’s expectations to guarantee their revenue (R = –0.11, Table 4).

Bergevoet *et al.* (2004) [10] had a considerably more consistent opinion. The “extra-source of income” profile producers (showing similarities with the LBE model) were more likely to declare that in their decision-making they take the environment into consideration, even if it lowers profit. The “large and modern farm” profile producers do not mention their will to adopt these initiatives.

#### Climatic hazard

Facing feed shortages due to unfavourable climatic conditions, producers tending toward GBI and LBE seem not to have the same way of thinking; GBI producers intend to buy high nutritional feed to balance the shortages (estimate = 0.22, Table 5) and LBE producers are going to decrease the number of cows (estimate = –0.25, Table 5) and ensure their feed autonomy (estimate = –0.17, Table 5).

### Current situation vs. ideal future farm

The current situation of dairy producers was compared to their preferred IFF (Table 1). Except for the type of workforce, quite high percentages of “unhappy” producers were observed for the farm characteristics, between 37 to 50%. This suggested that not all producers work as they would like to. The same comparison was not found in the literature, to our knowledge.

As dairy producers do not work in a way that they consider to be ideal, it is interesting to study the gaps to fill in order to reach their ideal system and so, amongst others, their needs. The study of the requirements to reach the IFF, including ways to meet these needs and the area of the needs, can inform the stakeholders of the dairy sector about what must be developed to evolve into IFF.

### Needs

#### Paths to formation

As way to improve their skills, producers wanting GBI tended to favour consultancy (estimate = 0.17, Table 6) and commercial companies (estimate = 0.16, Table 6) and not days of study on other farms (estimate = 0.082, Table 6), meanwhile producers wanting LBE supported this latter possibility (estimate = –0.088, Table 6), a network of pilot farms (estimate = –0.13, Table 6) and the associate, not market sector (estimate = –0.21, Table 6). Moreover, for help in technical choices, producers desiring LBE chose formation and study days (estimate = –0.15, Table 6) and producers’ technical groups to implement in the future (estimate = –0.20, Table 6). The choices presented confirm the will for a non-market way to learn for producers wanting LBE, contrary to producers wishing for GBI.

**Table 6.**
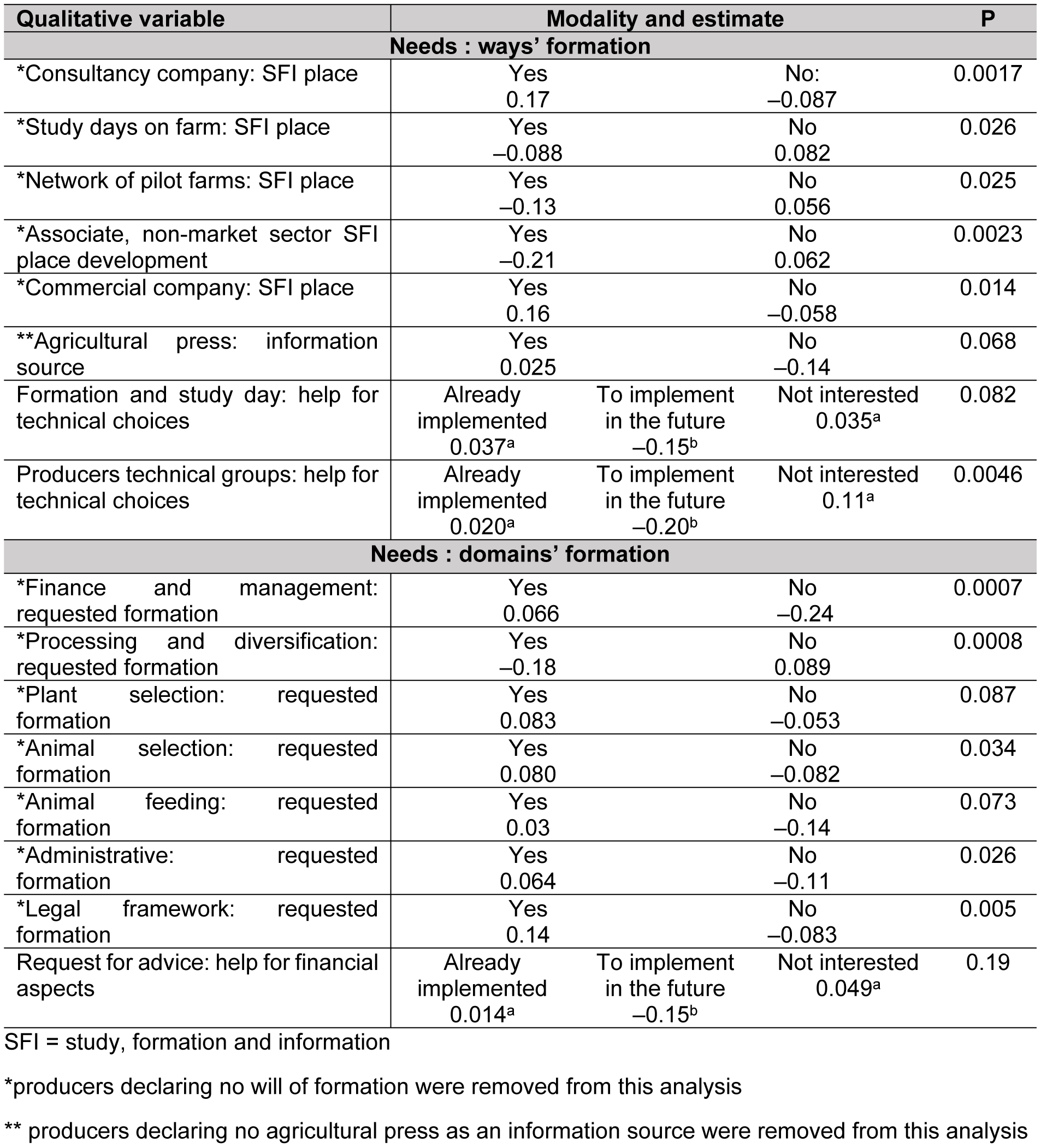
Value and level of significance of the difference in the ideal future farm gradient as a function of modalities of qualitative variables: formations

| Qualitative variable | Modality and estimate |  |  | P |
| --- | --- | --- | --- | --- |
| Needs : ways' formation |  |  |  |  |
| *Consultancy company: SFI place | Yes<br>0.17 | No:<br>−0.087 |  | 0.0017 |
| *Study days on farm: SFI place | Yes<br>−0.088 | No<br>0.082 |  | 0.026 |
| *Network of pilot farms: SFI place | Yes<br>−0.13 | No<br>0.056 |  | 0.025 |
| *Associate, non-market sector SFI place development | Yes<br>−0.21 | No<br>0.062 |  | 0.0023 |
| *Commercial company: SFI place | Yes<br>0.16 | No<br>−0.058 |  | 0.014 |
| **Agricultural press: information source | Yes<br>0.025 | No<br>−0.14 |  | 0.068 |
| Formation and study day: help for technical choices | Already implemented<br>0.037<sup>a</sup> | To implement in the future<br>−0.15<sup>b</sup> | Not interested<br>0.035<sup>a</sup> | 0.082 |
| Producers technical groups: help for technical choices | Already implemented<br>0.020<sup>a</sup> | To implement in the future<br>−0.20<sup>b</sup> | Not interested<br>0.11<sup>a</sup> | 0.0046 |
| Needs : domains' formation |  |  |  |  |
| *Finance and management: requested formation | Yes<br>0.066 | No<br>−0.24 |  | 0.0007 |
| *Processing and diversification: requested formation | Yes<br>−0.18 | No<br>0.089 |  | 0.0008 |
| *Plant selection: requested formation | Yes<br>0.083 | No<br>−0.053 |  | 0.087 |
| *Animal selection: requested formation | Yes<br>0.080 | No<br>−0.082 |  | 0.034 |
| *Animal feeding: requested formation | Yes<br>0.03 | No<br>−0.14 |  | 0.073 |
| *Administrative: requested formation | Yes<br>0.064 | No<br>−0.11 |  | 0.026 |
| *Legal framework: requested formation | Yes<br>0.14 | No<br>−0.083 |  | 0.005 |
| Request for advice: help for financial aspects | Already implemented<br>0.014<sup>a</sup> | To implement in the future<br>−0.15<sup>b</sup> | Not interested<br>0.049<sup>a</sup> | 0.19 |

SFI = study, formation and information
\*producers declaring no will of formation were removed from this analysis
\*\* producers declaring no agricultural press as an information source were removed from this analysis

As an information source, the agricultural press was commonly cited (N = 161, i.e., 78% of respondents), but producers desiring LBE tend to not want to inform themselves in this conventional way (estimate =–0.14, Table 6).

Producers wanting a GBI model tend to need more help to free them from their work in order to follow a formation (R = 0.21, Table 4)

#### Formation domains

The formation domains reflected the direction chosen by producers looking for LBE and the ways to reach it. They tend to want skills related to processing and diversification (estimate = –0.18, Table 6) and were likely to reject finance, management (estimate = –0.24, Table 6), administrative (estimate = –0.11, Table 6) and legal framework (estimate = –0.083, Table 6) skills. For financial aspects producers wanting LBE tend to favour requests for advice from experts rather than self-formation (estimate = –0.15, *P* _to implement *vs*. not interested_ = 0.12, Table 6). They do not choose animal feeding (estimate = –0.14, Table 6) and selection formations (estimates = –0.053; –0.082, Table 6). This could suggest the will of the producers not to change their way of management and the level of quality of their herd but the method of valorisation of their production.

In contrast, producers desiring GBI tend to want to continue enhancement of their vegetal and animal production (estimates = 0.083; 0.08, Table 6), to become more efficient and enhance their revenue. Moreover they are more interested in legal aspects (estimate = 0.14, Table 6). Expansion and complexification of the GBI model of dairy farms wished for by these producers could be an explanation. Bergevoet *et al.* (2004) [10] also observed a will to be well informed about the legislation for the “modern and large farm” profile. This is not noted in their profile, which is close to the LBE model.

Two kinds of formation were identified and preferred by producers wanting LBE or GBI models. Bergevoet *et al.* (2014) [10] observed the will to innovate for the two profiles closest to LBE and GBI profiles of this study. Verhees *et al.* (2018) [13] observed that formation was the most important resource for dairy producers. The present research differentiated the formation desired as a function of IFF. Dufour *et al.* (2007) [33] defined, through a survey of 15 dairy farmers, three conceptions of the work: difficult, organisational and passionate. The passionate approach was accompanied by the desire for new knowledge which was, as observed here, either to learn about genetic selection or about processing and marketing of products.

## Conclusions

In conclusion, the GBI tendency is two times more represented than the LBE tendency. Many reasons explain this choice of ideal farm. Past crises seem to cause farmers to desire the LBE model. A high workload seems to orientate respondents to the GBI model. The wish for the IFF is influenced by the current framework but is also a question of mentality. Production factors reached, breeds chosen for the herd, ways to react to factors external to the farm, consideration of diversification and alternative valorisation, regrouping, and mechanisation and robotisation describe the producers’ mentality and showed different relations with the IFF chosen. Moreover LBE and GBI producers may both have concern for environment, but the approach to act for the environment by LBE producers, through concern for ecosystem services, is clearly highlighted in this study. These producers found it important to answer to society’s expectations. Finally, as the current situation of farming is quite different to the ideal one, the needs for learning were studied and two types of customer appeared within dairy producers in relation to their formation. We conclude that two kinds of producers seem to appear, for different reasons, with different relations to the environment and asking for different formations.

## Acknowledgments

I want to thank the organising committee of Carrefour des Productions animales for the supply of the data.

